# From Scattered to Focused: Task-Dependent Connectivity in Honey Bees, with Midge Swarms and Bird Flocks

**DOI:** 10.1101/2025.11.21.689829

**Authors:** Ishriak Ahmed, Md. Saiful Islam, Imraan A. Faruque

## Abstract

Collective motions in biological organisms such as honey bees, midges, and birds exhibit remarkable coordination and adaptability, emerging from local interactions among individuals, including those with tight constraints on neural material. Understanding the underlying mechanisms behind the collective behavior of biological swarms informs the dynamics of decentralized systems, including engineered aerial swarms.

This study provides a systematic graph network extraction formulation applicable to physical domains trajectories including the practical effects of data dropouts, multiple agents, and multiple degrees of freedom. The analysis uses graph network modeling (in which nodes represent agents and edges denote interactions) to uncover the connectivity patterns governing swarm dynamics and quantify their differences. Three network identification methods, including sparse regression, Granger-causality, and crosscorrelation, are applied to determine the underlying graph from measured animal trajectories. An additional geometric anisotropy analysis used in bird flocks is also applied. The methods are applied to experimental recordings of honey bees involving scattered and focused “return-to-hive” behaviors, to midges in reproductive-related aggregations, and to flocking passerine birds. Across these studies, the number of agents a focal agent receives information from was quantified by analyzing the connectivity of identified interaction graphs.

The results reveal honey bees show significant differences in connectivity and slight differences in anisotropy between experiments involving scattered and focused “return-to-hive” behaviors. In particular, scattered honey bees showed causal relationships up to 10 neighbors (consistent with flocking jackdaw data), while focused honey bees showed connections with up to 2-3 agents, and midges showed slightly lower connectivity. Finally, cross-correlation analysis indicated honey bees exhibit distance-sensitivity in close proximity, while flocking birds exhibit a distance-dependent correlation decline. Together, these results provide evidence for task-dependent in-degree modulation in honey bees, and provide a systematic analysis formulation that highlights the variety of approaches to coordinated motion, including those with and without geometric anisotropy. These results inform the design of bioinspired networked systems.

## I. Introduction

The identification of graph networks in biological swarms is a crucial step toward uncovering the mechanisms that give rise to their complex collective behaviors. Swarming organisms such as honey bees, midges, and birds exhibit remarkable coordination and adaptability, despite each individual possessing limited sensing and computational capabilities. These sophisticated group-level patterns emerge from decentralized interactions among neighbors rather than from any global control. A central question in swarm biology is therefore determining the number of neighbors that influence a focal agent’s behavior and how this number varies across species, environmental conditions, and task demands. Representing such systems as graph networks, where nodes correspond to individuals and edges encode interaction strengths or information flow, provides a powerful framework for analyzing collective behavior. Graph-based system identification offers a principled way to reconstruct these interaction networks directly from trajectory data. In this study, we apply three different network estimation tools on experimental trajectories of honey bees, midges, and jackdaws, revealing their agent-to-agent connections. Such insights deepen our understanding of natural collective intelligence and inform the design of bioinspired engineered multi-agent systems.

## II. Previous work and background

### A. Collective motion and interaction mechanisms among species

A characteristic of biological systems is collective motion, which results from local interactions between individuals and produces coordinated group-level behaviors. Different species (fish, pigeons, goats, baboons) demonstrate diverse collective patterns such as polarization and group shape, reflecting species-specific communication mechanisms and adaptations to the environment [1]. Among insects, midge swarms have been extensively studied for the physics of collective behavior. Although midges display lower polarization than birds, their swarm structure indicates the presence of an effective elastic force that maintains group cohesion [2]. Environmental perturbations have been shown to induce velocity correlations within midge swarms [3], suggesting that these systems can dynamically respond to external stimuli while preserving internal organization. When midge swarms are subjected to external perturbations that mimic thermodynamic cycles, derived equations of state inspired by classical thermodynamics hold, suggesting that physical principles similar to ideal gas equations may also describe collective biological systems [4]. Pairwise interaction may play an important role in the emergent collective behaviors in midges. Time frequency analysis has shown two distinct modes of low frequency and high frequency pairwise interactions [5]. Interaction range in midges varied within 2-5 centimeters [6], consistent with their acoustic properties [7]. A similar acoustic range is found in mosquitoes [8], suggesting that an acoustic range can be a driver of swarm interaction in insects, visible through correlations in metric distances. Midge swarms tune their control parameter so that their correlation function can scale with increasing number of agents; therefore, an aggregation mechanism that supports maximum sustainable swarm size is plausible [9]. They also exhibit an inward velocity trend among insects flying at the edge of the swarm [10]. From disordered to highly aligned, movement behavior in marching locusts can arise solely from pairwise interactions and information transfer during flight, even without external disturbances [11]. These studies collectively highlight the diversity of mechanisms through which insects communicate, coordinate, and self-organize into cohesive swarms, demonstrating that complex collective behaviors can emerge from simple interaction rules.

Birds exhibit highly polarized and coherent flocking behavior, in contrast to the more disordered and diffuse motion typically observed in insect swarms. Pairwise or leader-follower behavior has been identified in bats and pigeons through spatial correlations [12, 13]. Birds that maintain pairwise interactions experience reduced workload when flying in pairs, and these interactions help flocks stay cohesive and navigate through obstacle fields more effectively [14, 15]. The interaction during flocking between starlings shows a topological nature, each bird connecting with approximately 7 neighbors [16, 17]. In spite of interaction with only a few neighbors, it has been shown that the range of spatial correlation scales with the linear size of the flock [18]. The degree of interaction may also vary in the same species. For example, within jackdaw flocks, paired birds interacted with fewer agents, and correlation lengths decreased in the presence of more paired birds [15]. While birds have been shown to maintain topological connections with a variable number of neighbors in different circumstances, it remains unclear how many individuals insects can simultaneously interact with, given their substantially limited neural processing capacity compared to avian species.

### B. Selective attention in insects

“Attention” often describes responsiveness to a subset of information at a given time, and “selective” attention capacity typically describes an animal’s capacity to respond to a task-relevant information subset. Previous works provide evidence of insects selectively responding to task-relevant stimuli, and their measured behavior shows that they disregarded other signals in their visual field [19], indicating some selective attention ability in visual sensing [20, 21]. Visual search experiments provide evidence that honey bees may attend to multiple stimuli sequentially rather than in a parallel method, more consistent with bumble bees or humans [22, 23]. Experiments finding longer decision times as the number of distractors increases support this distinction, suggesting that more demanding tasks reduce individuals’ capacity to attend to other events. Internal resource competition for limited working memory provides one possible mechanism for longer decision times with more complex discrimination tasks, and previous honey bee studies suggest a visual working memory interval of approximately 6–9 seconds [24]. Prior learning can also alter honey bees’ inherent preference for global visual information, enabling them to selectively attend to local features rather than global patterns within a stimulus [25].

Focused attention capacity is also described in predator–prey interactions in insects such as mantises and dragonflies. Neural recordings from dragonfly small target motion detection (STMD) neurons indicate the ability to alternate fixation among competing targets [26]. In multiple prey scenario experiments, mantises and dragonflies selectively track a single target while disregarding others [27, 26]. Similarly, fruit flies (*Drosophila melanogaster*) demonstrate the ability to direct their behavior toward stimuli located in specific regions of their visual field while ignoring other visual cues [28]. Across different insect species, a target’s spatial position can effectively guide orienting behavior in fruit flies, honey bees, dragonflies, and hoverflies [28, 29, 30, 31].

### C. Graph identification methods

Network inference, reconstructing inter-agent topology directly from observed behaviors, is an emerging field that extends principles of signal processing to graph-structured data [32, 33]. Examples of recent work on topology inference include Dong et al. [34], who learned graph Laplacians using a factor analysis model with Gaussian priors to ensure smooth signal variation, and Zhu et al. [35], who applied convex optimization to infer weighted, undirected graphs from consensus dynamics. For directed networks, Jiao et al. [36] developed an input-filtering approach to account for latent inputs, while Liu et al. [37] reconstructed robot interaction graphs directly from trajectory data. In visual data analysis, Ayazoglu et al. [38] combined causality and Lasso-based regression to identify directed relationships between entities. In biological contexts, topology inference often relies on neural or information-theoretic approaches [39], while engineered systems focus on explicit graph construction. In the present study, the graph identification problem is approached through multiple methods: least-squares error minimization with sparsity constraints, Granger causality analysis for directed connectivity, crosscorrelation for assessing pairwise coupling strength, and anisotropy for evaluating spatial organization. These integrated methods are applied to experimental biological flight data to reveal the underlying network structure and statistical properties of collective behavior.

### D. Contribution of the study

Despite substantial advances in characterizing coordination rules, significant gaps remain in our understanding of how biological systems achieve collective coordination, especially during high-bandwidth group tasks performed under strict neural and communication constraints. While interaction neighborhoods and network connectivities have been quantified in several large vertebrate groups, the limited neural capacity and comparatively coarse visual systems of insects suggest that their effective interaction neighborhoods may be far more restricted. Yet, no direct experimental measurement of neighborhood size in honey bee swarms has been carried out to date. In this study, we

1. developed network-inference methods based on observed velocities using sparse linear regression and Granger causality analysis, and cross correlation,
2. estimated graph-based interaction neighborhood sizes using biological trajectory data from insects and birds, and
3. identified evidence of task-dependent attention modulation in honey bees, reflected by a reduction in their effective interaction neighborhood during a focused task.

## III. Methods and Approach

### A. Animal datasets

In this work, flights of different biological species were considered, including insects (honey bees, midges) and birds. Two different kinds of honey bee flight data were used.

#### 1) Honey bee in scattered swarm

The dataset, sourced from Mahadeeswara and Srinivasan [40], features two processed videos of honey bees (*Apis Mellifera*) flying in dense swarms. Conducted in a lab setting, the hive entrance was temporarily blocked, causing returning foragers to form a dense “bee cloud” in front of the hive. Video recordings began ~ 30 seconds after entrance blocking and lasted up to 10 seconds before reopening the entrance. The videos were recorded at 200 fps with a resolution of 2048×1024 pixels. The two datasets used in this study had a recording length of 7.9 and 7.0 seconds.

#### 2) Honey bee in focused swarm

The focused task consisted of returning foraging honey bees (*Apis Mellifera*) in an outdoor task while they re-entered their hive through an actuated moving hive entrance in group conditions. Three-dimensional insect positions were optically tracked from a multi-camera system running at 20Hz in 5-second intervals [41]. Data collection occurred during daylight between 12 p.m. and 3 p.m. as approaching insects formed groups around the hive entrance.

#### 3) Midges in reproductive swarms

Dense swarming flying midge data was collected from Sinhuber et al. [42]. The data was collected by imaging swarms of *Chironomus riparius* midges from a self-sustaining laboratory colony housed in a (122cm)^3^ acrylic enclosure, designed for optimal optical access. Trajectories were captured using three synchronized Point Grey Flea3 cameras, capturing 8-bit greyscale images at a resolution of 1280×1024 pixels and a frame rate of 100 Hz. The swarms were illuminated with near-infrared LEDs, which were visible to the cameras and not to the midges to avoid disturbing the behavior. Each swarming event was recorded for 2 to 5 minutes. Our analysis is based on observations 15 and 17 with mean swarm sizes of 20 and 15 insects.

#### 4) Jackdaw flocks

The flocking bird dataset consisted of the Ling et al. [15] dataset, which contains 3D trajectory data of wild jackdaws (*Corvus monedula*) while their movements were optically tracked using four tripod-mounted Basler Ace cameras (2048×2048 pixel resolution, 5.5 µm pixel size), recording at up to 90 frames per second.

Each dataset contained trajectories of multiple agents with unique identification numbers. The data were segmented into temporal windows for graph identification of interaction structures. Incomplete trajectories within segments required the removal of some agents. Smaller windows reduce data loss but may lead to underdetermined least-squares problems with insufficient equations. Thus, the window length was optimized to balance data completeness and solvability by retaining the maximum number of valid agents while ensuring stable parameter estimation. Agent removal due to data dropouts is an inherent observability limitation. A fixed window length was used for each video sequence.

### B. Inter-agent connection analysis

Inter-agent connectivity analysis consisted of building an estimate of the agent-to-agent graph representing the influence network (for which three complementary methods were used in parallel to provide robustness to method), then in-degree determination applied to each method, computing correlation distance, and an anisotropy analysis previously used on passerine flocks.

#### 1) Network identification by sparse linear regression

Given measured flight trajectories of animals, the objective is to infer inter-dependencies among agents. Let, 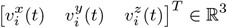, be the velocity of agent *i* at timestep *t*, in the presence of *N* total agents. A one-step-ahead update law of each agent can be written as

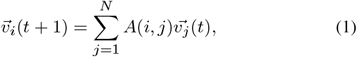

where *A*(*i, j*) represents the interaction weight quantifying the influence of agent j on the velocity update of agent i. The inter-agent weights can be collected into the matrix A ∈ ℝ^*N×N*^. The objective is to estimate this matrix from the trajectory data and interpret its entries as a representation of the underlying interaction graph among the agents. To estimate A, a least squares regression problem is formulated. For each timestep *t* and dimension *x*, the update rule can be expressed as

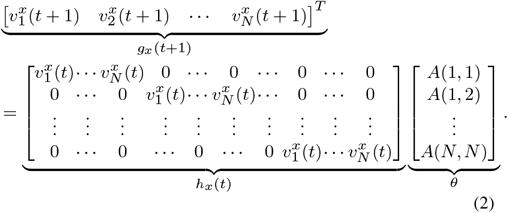

Now, all measurements can be concatenated vertically over the three dimensions *x, y, z* and over all time steps *t* = 1, 2, …, *t*_*f*_ to form a single linear regression problem as

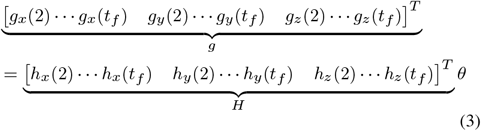

This format is suitable for solution via least squares regression, where the number of datapoints in the observation set is *n*_*d*_ = 3N(*t*_*f*_ −1) with 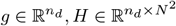 and 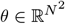. To uniquely identify θ, the number of observations must satisfy *n*_*d*_ ≥ *N* ^2^, which yields a constraint on the minimum timesteps

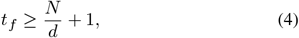

where d = 3 is the dimension of the observed state (3D velocity).

If no structural constraints are applied on *θ* or equivalently the matrix *A*, the solution can be found by solving the unconstrained least squares optimization problem

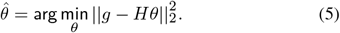

The corresponding identified interaction matrix is denoted by *Â*_*U*_. Since numerical solvers are used to compute *Â*_*U*_, the entries of *Â*_*U*_ are non-zero due to the absence of regularization, even if some interactions are negligible or non-existent. A **sparse** identification technique promotes automatic elimination of negligible interaction terms by encouraging many entries of θ to become zero. To achieve this, an *ℓ*_1_ regularization term is added to the objective function, resulting in the optimization problem

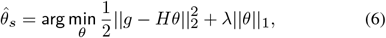

where λ is the regularization parameter controlling the trade-off between model fit and sparsity. This formulation (e.g., the MATLAB `>>lasso` function) penalizes small-magnitude coefficients to drive many of them to zero. The resulting matrix *Â*_*S*_ is therefore sparse, maximizing interpretability of the agent-to-agent interaction representation.

#### 2) Graph based on Granger causality

Granger causality [43] is a statistical framework for detecting causal relationships between time series data. The fundamental principle is that if the past values of one variable improve the prediction of another variable beyond what can be achieved using the latter variable’s own past, then the first variable is said to *Granger-cause* the second. This method is capable of accounting for shared underlying influences among variables. Initially developed in econometrics, Granger causality has found widespread use in neuroscience, biology, and other fields [44]. In the context of biological swarm trajectories, Granger causality thus provides a directed measure of inter-agent information transfer, quantifying how the motion of one agent influences or predicts that of another. For hypothesis testing in this study, a first-order vector autoregression model (VAR) was employed. The method is briefly outlined below. Similar to the unconstrained identification approach described with the model structure in Eq. (1), a first-order unrestricted vector autoregressive (VAR) model can be formulated as

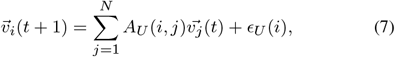

where ϵ_*U*_ (i) is the error residual in the unrestricted model. A reduced VAR model, omitting the contribution of agent k, can be written as

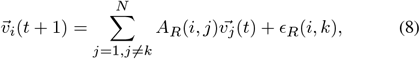

where *A*_*R*_ represents the restricted weight matrix. Granger causality statistically quantifies the extent to which the unrestricted model in Eq. (7) provides a better fit than that in Eq. (8). If the error ϵ_*R*_(i, k) is significantly larger than ϵ_*U*_ (i) in a statistical sense, it implies that agent k’s contribution to the fitted model can not be overlooked and therefore a causal relation between agent i and k is inferred. Granger causality evaluates the null hypothesis *H*_0_ : *A*_*U*_ (i, k) = 0 using an F-test to determine whether agent k has a statistically significant predictive influence on agent i. A significance threshold of p < 0.05 is adopted to reject the null hypothesis, indicating a significant causal effect. The resulting p-values for all agent pairs are assembled into a matrix G, which represents the pairwise causal connectivity structure of the system. The computations of Granger causality statistics and corresponding p-values were performed using the multivariate Granger causality (MVGC) toolbox [44].

#### 3) Graph based on cross-correlation

Cross-correlation between two time signals quantifies their similarity as a function of temporal alignment, using the normalized dot product between the signals. To determine the degree of correlation between different agents, a cross-correlation analysis was performed on their positional time-series trajectories, providing a measure of how similarly pairs of agents move over the time window. From time histories of agents, let 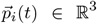 be the position vector of agent *i* at timesteps *t* = 1, 2 · · · *t*_*f*_. Considering a time-delay *τ*, the cross-correlation between two agents *i* and *j* is defined as

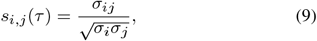

where

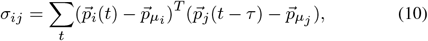

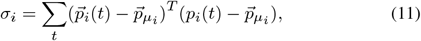

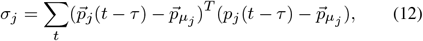

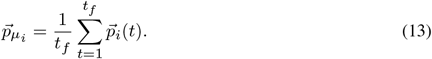

The correlation weight between agent *i* and *j* is defined as

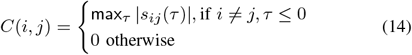

#### 4) In-degree determination

In a directed graph, the *in-degree* of a node represents the number of directed edges entering that node. In the context of the multi-agent system considered here, each agent is modeled as a node, and the interaction strengths between agents are represented as weighted edges. The in-degree thus quantifies the number of neighboring agents that exert influence on a given agent, or equivalently, the number of agents to which it allocates attention. This in-degree quantification consists of transforming weight matrices into a binary connectivity matrix B in which each entry *b*_*ij*_ = 1 indicates the presence of a directed connection from agent *j* to agent *i*, and *b*_*ij*_ = 0 otherwise. In-degree is found by summing up the rows of B using

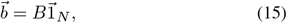

where 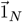 is a column vector of N ones. The *k*^*th*^ row of 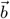 provides the in-degree of agent *k*.

The identified interaction weights obtained from the sparse regression, Granger causality, and cross-correlation analyses are processed to construct a binary connectivity matrix

##### a) From sparse identification

For sparse identification, the corresponding binary connectivity matrix *B*_*S*_ is given by

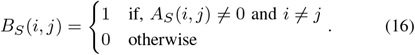

##### b) From cross-correlation

To determine statistically significant connectivity from cross-correlation between agent signals, it is necessary to account for the effect of finite signal length. Two independent white-noise sequences of length t_*f*_ exhibit a 95% confidence interval for their cross-correlation values given by 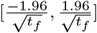 [45]. Any observed cross-correlation value *C*(*i, j*) beyond this interval was considered significant, indicating a non-random correlation between agents *i* and *j*. Two agents are connected in a binary connectivity matrix *B*_*C*_ such that

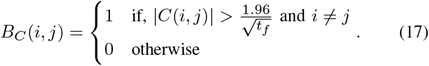

where *t*_*f*_ is the number of time steps.

##### c) From Granger causality

Based on the results of the Granger causality analysis, the corresponding binary connectivity matrix *B*_*G*_ was constructed by identifying statistically significant interactions at a significance level of *p* < 0.05, such that

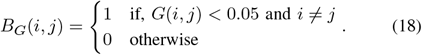

The final in-degree of an agent can be determined by summing up the binary matrices B_*S*_, B*X*, or B*G* row-wise.

#### 5) Correlation distance

To examine how the identified inter-agent connection weights vary as a function of metric distance, the mean inter-agent distance over a data segment was defined as

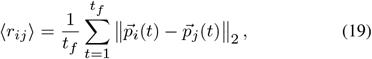

where 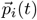 and 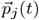 denote the positions of agents *i* and *j* at time step t, respectively, and *t*_*f*_ is the total number of time steps in the trial. This average Euclidean distance ⟨r⟩ serves as a measure of spatial proximity, which is then related to the corresponding inter-agent correlation weights.

To analyze the variation of cross correlation as a function of the average inter-agent distance ⟨*r*⟩, the cross correlation weights in *C* were grouped into 50 bins spanning the full range of observed inter-agent distance. For each bin, the number of data points was recorded, and the corresponding mean ⟨*C*⟩ and range of the edge weights were computed to characterize the distribution of correlation strength as a function of distance.

#### 6) Anisotropy analysis

Empirical investigations of orientation patterns in starling flocks have revealed a distinctly anisotropic spatial organization [46]. This anisotropy implies that individuals preferentially position themselves relative to one another along specific directions defined by their global velocity vector. To quantify the strength of this directional bias with a single scalar measure, the *anisotropy factor*, denoted by γ was introduced [46]. The computational procedure for evaluating this metric follows the methodology outlined in [47]

Given a certain group, at timestep t, consider the set of all *first* nearest neighbor directions 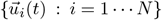 where *N* is the number of individuals. For each one of these vectors,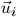 the following matrix

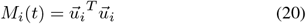

can be built, which works as a projection matrix. The effect of multiplying by this matrix with an arbitrary vector 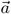 is to produce a new vector in the same direction as 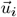 with a magnitude of the projection of 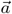 along 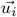. The average of all projection matrices over all agents

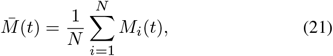

produces a matrix that projects along the average direction of the *first* nearest neighbor. The eigenvector 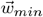 corresponding to the smallest eigenvalue indicates the direction along which there is the smallest number of nearest neighbor vectors; therefore, it indicates the average direction of *minimal crowding*. The corresponding anisotropy factor γ_min_ is defined by

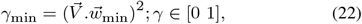

where 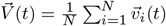 is the group average velocity.

Similarly, γ_max_ is defined as the component corresponding to the eigenvector associated with the maximum eigenvalue, representing the direction of maximal crowding. Geometrically, an average value of γ = 1/3 indicates an isotropic, fully decorrelated spatial distribution, whereas deviations from this value signify the presence of anisotropic spatial structure. Since there is no established evidence that insects exhibit a consistent preference for either minimal or maximal crowding directions, anisotropy is examined in both orientations. While the example above defines anisotropy based on the first nearest neighbor, the same formulation can be extended to the second, third, and higher-order neighbors.

In swarm systems, border agents experience noticeably different spatial conditions compared to those within the interior. Unlike inner agents, which are surrounded, border agents have few or no neighbors on one side, leading to asymmetric local environments. Consequently, it is essential to account for these border effects when computing anisotropy. To identify border agents, a bounding surface was generated to enclose the set of three-dimensional positions representing the swarm. This was achieved by constructing a triangulated boundary around the agent positions. The boundary surface was then adaptively shrunk toward the interior of the convex hull to ensure that most peripheral agents were correctly classified as border members. This study used the MATLAB >>boundary function with a shrink factor of 0.5 to provide a balance between overestimating and underestimating the boundary region.

## IV. Results and discussion

### A. Spatial anisotropy in agents

Anisotropy analysis reveals the geometric organization of the relative positions among agents with respect to their average direction of motion. This approach has been widely applied in studies of bird flocks to identify positional asymmetries and preferential alignment patterns, thereby uncovering the underlying inter-agent connectivity structure. Here, the same analysis was extended to insect groups to investigate whether any inherent geometric regularities exist. The results are presented in Fig. 2.

**Fig. 1:**
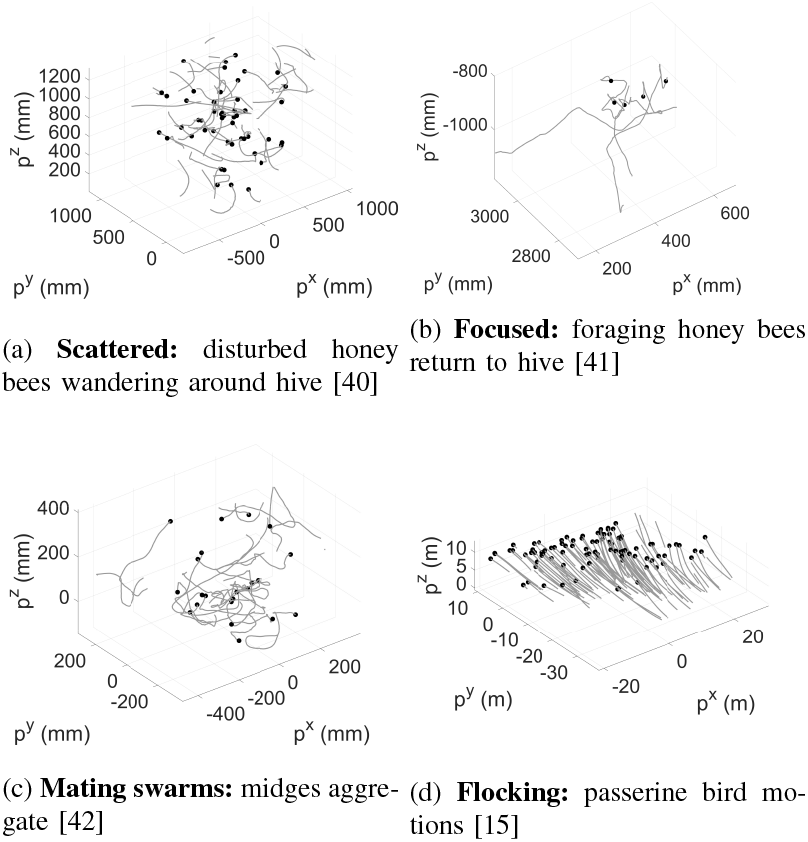
This study analyzed data from four biological flight experiments: (a) scattered honey bees in a return-to-hive behavior, (b)a focused foraging return-to-moving hive task, (c) stationary mating midge swarm aggregations, and (d) flocking passerine birds, providing insight into organism and task-specific differences.

**Fig. 2:**
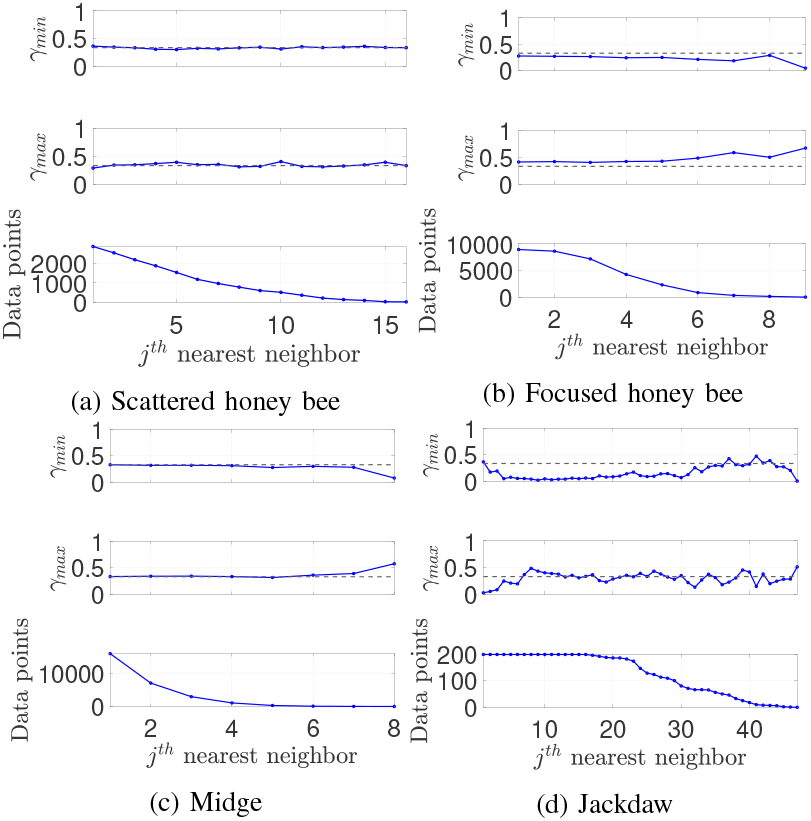
Anisotropy analysis results in biological datasets. The dashed line at γ = 0.3 shows the theoretical isotropic value. All datasets were processed through border correction and Hanisch correction, except for focused honey bees. Scattered honey bees and midges (a,c) showed no anisotropy. Focused honey bees (b) showed slight anisotropy in both minimal and maximal crowding direction. γmin being slightly less than 0.3 suggests the tendency of minimal crowding sideways, and γmax slightly greater than 0.3 indicates a slight line-forming tendency, consistent with the fact that they are all trying to enter an entrance one after another. Jackdaws (d) showed maximal crowding tendency perpendicular to the group average velocity in the considered dataset.

In this analysis, both γ_min_ and γ_max_, corresponding to the minimal and maximal crowding directions, were considered. An anisotropy value close to one indicates that the corresponding crowding direction is aligned with the overall direction of motion, whereas a value approaching zero signifies a perpendicular configuration—implying that the neighboring agents are positioned perpendicular to the average direction of motion. The theoretical baseline value of anisotropy is approximately 0.3; deviations from this value indicate a directional bias or spatial preference within the group structure.

#### a) Scattered insects showed no anisotropy

For both scattered honey bees and midges, the anisotropy analysis revealed no significant directional bias, with γ ≈ 0.3 across the analyzed datasets. This indicates an absence of preferential alignment between the overall group motion and the spatial arrangement of individuals. In other words, the collective dynamics of these insect groups do not necessitate any geometric pattern causing asymmetry in maximal or minimal crowding directions. Instead, their coordination appears to emerge from local interaction rules rather than from stable spatial organization. Dynamical systems approaches, as presented in this study, therefore provide a more effective framework for identifying such interaction mechanisms than purely geometric analyses.

By contrast, jackdaws exhibited a slight anisotropy for the maximal crowding direction involving approximately eight neighbors, in agreement with previous observations in avian systems, suggesting the tendency to align in the perpendicular direction with respect to the direction of motion.

#### b) Focused honey bees showed slight anisotropy

Focused honey bees exhibited a slightly lower anisotropy value than 0.3 in the minimal crowding direction and a slight increase in the maximal crowding direction. This suggests a weak tendency for individuals to align parallel to the average direction of motion. Such an effect is not unexpected, as the focused honey bees were observed sequentially entering the hive entrance, a task-driven behavior that naturally introduces mild geometric asymmetry in their spatial organization. However, anisotropy measurements can be sensitive to border effects, which, if not properly accounted for, may produce misleading results [47]. The Hanisch criterion [48] also recommends excluding agents located near or beyond the boundary of the swarm to avoid edge-induced bias. In the focused honey bee dataset, the presence of a maximum of 10 agents significantly limits the ability to define a meaningful border region. For this reason, border corrections were not applied, as doing so would have eliminated nearly all agents under the Hanisch filtering condition. Anisotropy analysis is therefore more informative for large groups with well-defined spatial boundaries, in contrast to the relatively ill-defined boundaries found in insect swarms.

### B. In-degree of agents

The in-degree of each identified binary connectivity matrix quantifies the number of neighboring agents that an individual interacts with, revealing the network structure. The three estimation methods employed *Sparse identification, Granger causality*, and *Cross-correlation* play complementary roles in interpretation.

The *Sparse identification* approach encourages numerical parsimony by seeking the smallest possible number of non-zero entries in the binary adjacency matrix, which makes it effective in isolating dominant or essential connections. However, this method may overestimate the number of weak links with noisy data by assigning near-zero weights to functionally insignificant edges, which nonetheless appear as connections in the binary adjacency matrix. It can also underestimate connections in the cases of highly synchronized behaviors, where assigning weight to one agent with similar velocity negates the necessity to assign weight to the other, by treating multiple correlated agents as redundant contributors to the focal agent’s behavior. Given these limitations, the peak of the resulting in-degree probability distribution can be interpreted as a reliable approximation of the underlying interaction network.

In contrast, the *Granger causality* analysis is grounded in statistical inference. It determines whether the time series of one agent provides significant predictive information about another agent’s future state. This property makes it suitable for finding directional dependencies and temporal causality, offering an interpretable measure of influence strength between agents. Granger causality explicitly tests for cross-agent effects, since every time series is trivially caused by itself. Granger causality may provide one of the closer approximations to an actual agent-to-agent connection, as it captures statistically significant predictive influences between time series. Since it identifies a connection as meaningful only when the agent’s own history cannot explain the effect, it imposes a particularly strict criterion. For this reason, the in-degree at 95% cumulative fraction of the data was used as an estimate of the maximal number of possible connections present within the dataset after accounting for outliers by filtering out potential outliers and numerically induced spurious links.

The positional cross-correlation weights measure mutual synchrony rather than causal links. Two correlated signals do not imply causation, because that correlation might arise from a shared external factor. Even in the absence of correlation, causal dependencies may still exist, masked by noise or nonlinearities in the system. Therefore, cross-correlation gives a measure of how visually coherent the motion appears, with highly polarized motions having high cross-correlation values.

With these in mind, this study highlights four noteworthy connectivity observations across the datasets:

#### a) Scattered honey bees showed a causal relationship up to 10 neighbors

According to the Granger causality weights shown in Fig. 3, honey bees showed a causal relationship to approximately 10 neighbors when considering the 95% cumulative fraction of the data. The *Sparse identification* method identified about 8 effective neighbors with the peak in its probability density function (PDF). This peak value in the sparse identification analysis closely aligns with the value at 95% cumulative distribution function (CDF) threshold from the Granger causality analysis, suggesting consistent structural connectivity estimates between the two methods. The *Cross-correlation* weights indicate synchrony extending from at least five agents up to 33 within the 95% cumulative fraction with a peak value at ~20 agents, showing the presence of group-level synchrony.

**Fig. 3:**
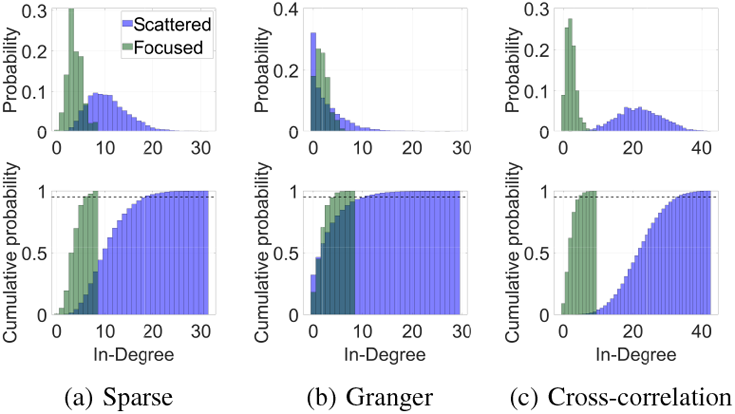
Inter-agent connectivity among honey bees quantified using three approaches: sparse identification, Granger causality, and positional cross-correlation analysis. In the sparse identification results, the *scattered* honey bee dataset exhibited a peak connectivity of approximately eight neighbors, whereas the *focused* dataset peaked at around three neighbors. The dashed line in the connectivity distribution represents the 95% cumulative fraction of data. At this threshold, the scattered dataset indicated ten connections when evaluated using Granger causality. Significant positional cross-correlations were observed with up to 33 neighboring agents. For the *focused* honey bee dataset, all three methods consistently indicated a lower in-degree of approximately three to four neighbors, both in terms of dynamic connections and cross-correlation measures.

#### b) Focused honey bees connected with 2-3 agents

Applying the Granger causality method with a 95% cumulative threshold, the focused honey bees exhibited directed interactions with at most four neighboring agents. The distribution of connections peaked at 1-2 agents, while the cross-correlation analysis indicated synchrony with approximately 1-3 agents. This pattern is consistent with the dyadic interaction behavior reported in previous studies, where a proportional–integral–derivative (PID) control rule successfully described the pairwise motion dynamics of agents in the same dataset [49]. Across the three analysis methods (*Sparse identification, Granger causality*, and *Cross-correlation*), the results remained generally consistent for this dataset. The focused honey bee dataset included at most ten agents flying together with two differences in characteristics from the scattered dataset: the smaller number of agents reduces statistical diversity, and the agents perform different behavioral tasks, leading to inherently distinct coordination dynamics. These factors together may account for observed differences between the results.

#### c) Midge swarms exhibited slightly lower connectivity than honey bees

Across all methods, as shown in Fig. 4, the midges demonstrated overall lower connectivity compared to the scattered honey bees. In the *sparse identification* analysis, the in-degree distribution peaked at four neighbors and reached up to six within the 95% cumulative fraction. The *Granger causality* analysis similarly indicated a maximum of up to six significant connections per agent, with approximately 45% of agents exhibiting zero in-degree. *Crosscorrelation* in-degree peaked near six, suggesting that interactions may be organized into multiple clusters whose influence does not necessarily span the entire swarm at once. However, these results may be lightly shaped by the smaller number of individuals in the midge dataset, which provides fewer possible connections than the honey bee recordings.

**Fig. 4:**
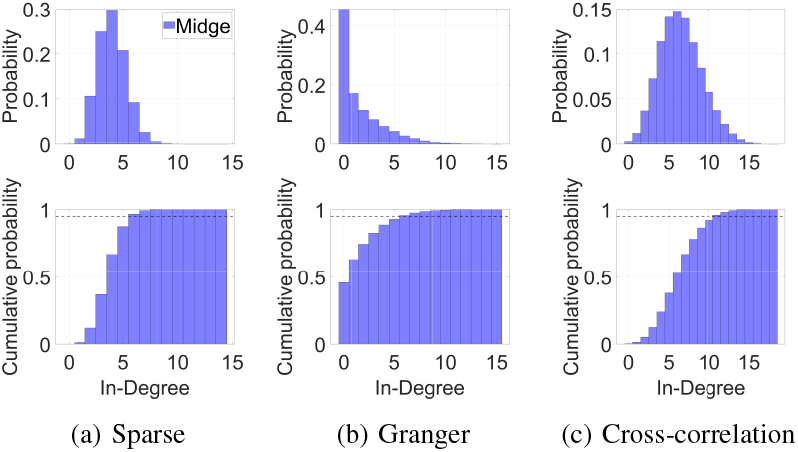
Midges exhibit lower overall connectivity, with in-degree distributions peaking at approximately four neighbors in sparse identification and six neighbors based on the 95% cutoff in Granger causality, with 45% midges showing no connectivity, and a similar peak of six in cross-correlation analyses.

#### d) Flocking jackdaws showed causal relations up to 10 agents

Birds exhibit more visually coherent and polarized motion compared to insects. As shown in Fig. 5, based on the Granger causality analysis, the 95% cumulative filter indicates that each jackdaw maintains causal relationships with up to ten neighbors. This observation aligns with earlier studies on starling flocks [16], which reported that birds tend to follow a *topological* interaction rule involving approximately seven to eight nearest neighbors, rather than a purely metric one, and the number of effective connections scales with flock size. In the present dataset, the original study [15] revealed the presence of *pairing behavior*, characterized by anomalously close flight between two individuals. Such a pairing could locally reduce the effective causal degree to one, as the dynamics of each pair become strongly coupled. However, when birds maintain cohesion with the larger flock, additional connections beyond the pair are required, raising the causal degree back toward the observed values. The *Sparse identification* method, in contrast, yielded a peak in-degree of three and a maximum of four connections, which is consistent with its design principle of enforcing parsimony. Because bird trajectories are highly polarized and often redundant in velocity space, the algorithm naturally discards correlated neighbors as superfluous predictors. This strong alignment is further reflected in the *Cross-correlation* results, where at least ~ 20 birds are found to be mutually correlated in a time window, highlighting the high degree of global synchrony within the flock.

**Fig. 5:**
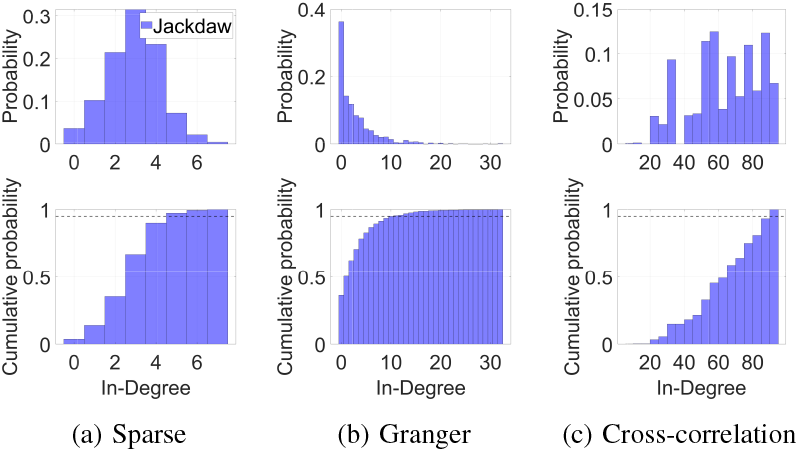
Jackdaws showed up to ten causal neighbors under the 95% Granger cutoff, while Sparse identification peaked at three strong connections, and high synchrony was evident in the cross-correlation in-degree.

### C. Variability of cross-correlation weights

In this analysis, the relationship between the cross-correlation weights and the inter-agent distances is analyzed, which reveals the tendency of distance-dependent synchrony. The plots in Fig. 6 represent the time-averaged inter-agent distance, ⟨*r*⟩, versus the average cross-correlation, ⟨*C*⟩, computed within discrete distance bins. Hence, the results illustrate an overall averaged trend rather than instantaneous pairwise correlations.

**Fig. 6:**
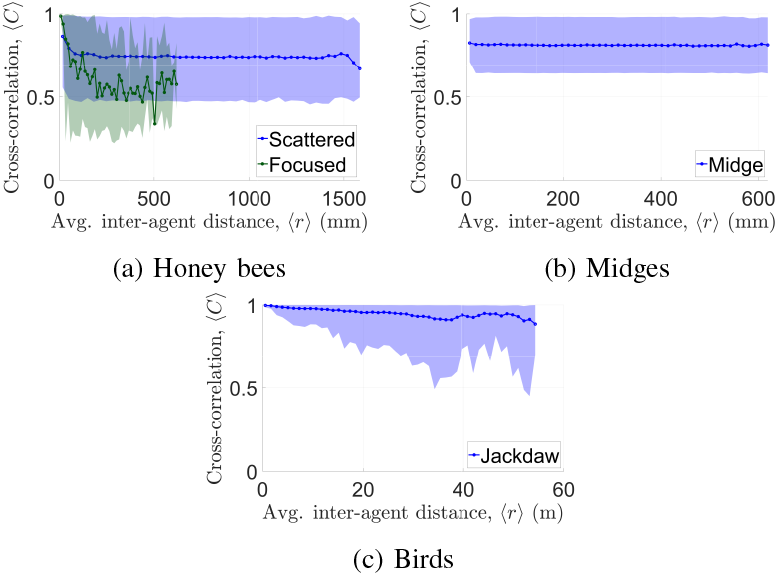
Dependence of Cross-correlation on Inter-agent Distance. Statistically significant cross-correlation is found for all species over the observed inter-agent distances. The mean value of ⟨*C*⟩ and its ranges are shown. In both focused and scattered honey bees, ⟨*C*⟩ decreased with increasing inter-agent distance up to ~100 mm, suggesting the presence of a decision zone. Cross-correlation in midges remained relatively uniform over inter-agent distances. Jackdaws showed a trend of decreasing ⟨*C*⟩ with increased distance while maintaining higher average values than insects.

Across different species, differences in correlation strength were observed. Birds exhibited the highest levels of polarization and correlation overall, a result not unexpected given the enhanced coordination mechanisms and sensory-motor control typical of vertebrate flocking systems compared to insect groups. Correlation levels in both scattered and focused honey bee flights varied across datasets, indicating a dependence on behavioral context. The following observations can be made regarding how spatial proximity influences the degree of collective synchrony across different species.

#### a) Honey bees showed distance sensitivity to cross-correlation in close proximity

In both the focused and scattered groups, an increased trend in average cross-correlation was observed at inter-agent distances below approximately 10 cm. Our previous study on paired honey bees indicated that individuals enter a *decision zone* within this range, after which they may either engage in pair formation or disengage [49]. The elevated cross-correlation at short distances is therefore likely associated with the increased number of dyadically interacting agents. This trend was consistently found in both focused and scattered honey bees. Overall, the focused honey bees exhibited lower mean correlation values than scattered ones at larger inter-agent separations but showed higher correlation in close proximity. This behavior can be interpreted in the context of task-driven motion: focused honey bees were observed approaching a nest entrance or target. At greater distances, their trajectories are dominated by goal-oriented flight rather than neighbor following. Upon entering the decision zone near the target, individuals begin to coordinate or pair with nearby agents, leading to a sharp rise in local synchrony and coherent motion.

#### b) Flocking birds exhibited a distance-dependent decline in correlation

The average cross-correlation ⟨*C*⟩ decreased almost linearly with increasing inter-agent distance in bird flocks. This dependence indicates that spatial proximity plays a strong role in maintaining coordinated motion among birds: agents that are physically closer exhibit more synchronized trajectories, whereas distant ones behave more independently. In contrast, correlation in midges remained relatively uniform across different separations.

### D. Perspectives on results

From a selective attention perspective, this study’s findings are some of the first evidence suggesting that individual honey bees can modulate the number of neighbors they attend to depending on the behavioral context. Rather than maintaining a fixed interaction neighborhood, agents may dynamically allocate their resources to the most relevant neighbors, enabling efficient coordination while minimizing processing load. Previous work indicated insects may be capable of filtering information from their visual field and prioritizing specific stimuli over others [28]. Behavioral patterns of honey bees indicate a tendency to process visual cues sequentially rather than in parallel [22], consistent with an interpretation that their attentional systems can focus on one neighbor or object at a time before shifting to the next. This sequential attention mechanism may provide an efficient strategy for insects with limited neural capacity, allowing them to selectively track only the most behaviorally relevant neighbor when the task demands focused, time-sensitive performance, such as homing toward the hive.

From a cognitive limitation perspective, the modulation of neighborhood size observed in our results may reflect a form of context-sensitive information filtering: bees appear capable of expanding or contracting their effective interaction set depending on whether they are navigating freely or performing a goal-directed task. In humans, the “magic number seven” hypothesis [50] identifies a limit on the number of items that can be simultaneously tracked or stored in working memory. Interestingly, similar numerical bounds have emerged in the analysis of topological interactions in bird flocks [16]. This parallel raises the possibility that such limits may be a general feature of biological systems operating under cognitive, sensory, and physical embodiment constraints.

From a dynamical systems perspective, in-degree variation affects the rate of information propagation in multi-agent networks. Simulations show that increasing the number of neighbors accelerates consensus formation, and that these gains plateau around ten neighbors, a regime nearly equivalent in performance to fully connected networks [51]. Empirical studies in midge swarms likewise indicate that key statistical descriptors of collective motion saturate once neighborhood sizes reach approximately an order of ten individuals [52]. These results suggest that collective systems may not benefit substantially from interaction in neighborhoods larger than this scale. The dependence of cross-correlation on distance in honey bees suggests that topological and metric frameworks are not mutually exclusive; rather, they can serve as complementary mechanisms that together shape the structure and stability of collective motion [53]. Together, these perspectives highlight that the regulation of neighborhood size is a multifaceted phenomenon shaped by perceptual, cognitive, and dynamical constraints inherent to biological swarms.

## V. Summary and conclusion

In this study, the aerial multi-agent connectivity structures and collective motion characteristics of three distinct biological systems—honey bees, midges, and jackdaws were analyzed using complementary analytical frameworks, including Sparse identification, Granger causality, and Cross-correlation. These approaches together provided a multifaceted view of how local interactions and global coordination emerge across different animals and behavioral contexts. Anisotropy analysis suggested that scattered insects, both honey bees and midges, showed no directional bias in their spatial arrangement. Focused honey bees exhibited a slight asymmetry, likely reflecting sequential alignment during entry into the hive entrance. In contrast, bird flocks show anisotropy involving approximately eight neighbors, aligning with established observations of directional structuring in avian groups.

Scattered honey bees exhibited causal connections extending up to approximately ten neighbors, whereas focused honey bees, which were engaged in goal-oriented flight toward a hive entrance, showed coupling with only two to three agents. This reduction in effective connectivity reflects a change from group-level coordination to task-driven motion dominated by individual or dyadic interactions, a significant transition in an invertebrate with limited neural capacity. Midge swarms displayed slightly lower connectivity than honey bees. The jackdaw flocks exhibited causal interactions involving up to ten neighbors, consistent with prior observations in other bird species such as starlings, while also showing a noticeable tendency to concentrate stronger connections on just a few (1–3) neighbors.

Jackdaws exhibited the highest correlation levels overall, reflecting the strong alignment and coordination typical of vertebrate flocks, and also exhibited an approximately linear decay of average correlation with increasing distance, indicating modulation of alignment strength across metric scales. Correlation level varied over data sets due to behavioral differences in honey bees. They displayed clear distance sensitivity, with increased correlation at distances below 10 cm, consistent with the “decision zone” observed in previous pair-bonding studies.

Overall, these findings demonstrate that the degree of connectivity, correlation, and spatial organization patterns varied with both species and behavioral context. Insects show emergent motion in spite of the absence of sustained geometric organization, whereas birds exhibit topologically structured and distance-dependent coordination. The complementary use of sparse identification, Granger causality, and cross-correlation thus provides complementary frameworks for uncovering the multiscale organization of collective motion across biological systems.

## Acknowledgments

This work was supported in part by ONR Young Investigator Award N00014-19-1-2216.

